# The preoptic Kisspeptin/nNOS/GnRH (KiNG) neuronal network regulates rhythmic LH release through a dual activation-inhibition mechanism

**DOI:** 10.1101/2024.01.15.575688

**Authors:** Virginia Delli, Charles-Antoine Seux, Julien Dehame, Sooraj Nair, Tori Lhomme, Konstantina Chachlaki

## Abstract

Gonadotropin-releasing hormone (GnRH) neurons are the final common target of a complex network of cells cooperating for the central control of reproduction. The balance between excitatory and inhibitory transsynaptic and non-synaptic inputs is crucial for the maintenance of the GnRH rhythms: the pulse and the surge. The precise mechanisms behind this remain under debate. In this work, we challenge the hypothesis that excitatory and inhibitory inputs from kisspeptin and neuronal nitric oxide (NO) synthase (nNOS)-expressing neurons orchestrates GnRH release, in a microcircuit that we call the Kisspeptin/nNOS/GnRH (KiNG) neuronal network. Our work specifically focuses on the role of nNOS neurons located in the organum vasculosum laminae terminalis (OV) and the median preoptic nucleus (MePO). nNOS and kisspeptin neurons interact anatomically and functionally, with the kisspeptin receptor (Kiss1r) being differentially regulated in nNOS-expressing neurons across the female estrous cycle. Using a novel viral tool allowing for the measurement of NO/cGMP levels with exquisite sensitivity, we demonstrate that kisspeptin is able to induce NO-dependent cGMP production in the OV/MePO, including in GnRH neurons *in vivo*. Using electrophysiological, genetic, chemogenetic and pharmacologic approaches, we reveal that NO production from nNOS neurons in the OV/MePO is needed to fine-tune the GnRH/LH response to kisspeptin, and specifically to turn off GnRH release, thus generating pulses. Our findings provide valuable insights into the tripartite KiNG neuronal network governing the regulation of the GnRH/LH pulse and surge.

## INTRODUCTION

Gonadotropin-releasing hormone (GnRH) neurons in the hypothalamus orchestrate reproduction. While their cell bodies are mainly distributed in the preoptic region, their nerve terminals, which lie in the median eminence of the hypothalamus, release the GnRH decapeptide into the pituitary portal blood system (Herbison, 2015; Moenter, 2017). Blood-borne GnRH acts on gonadotrophs in the anterior pituitary, where it stimulates the synthesis and secretion of the gonadotropins, luteinizing hormone (LH) and follicle-stimulating hormone (FSH). The release pattern of GnRH includes both pulse and surge phases in females but only pulses in males. GnRH rhythmicity arises from the integration of a complex array of excitatory and inhibitory inputs coming from both the central nervous system, e.g., kisspeptin and nitric oxide (NO), and peripheral organs (e.g., estrogen and testosterone from the gonads). The correct balance of these signals is indispensable for the timely and appropriate pattern of release of LH and FSH, key to the establishment and proper function of the hypothalamic-pituitary-gonadal (HPG) axis.

The stimulatory effect of kisspeptin-kisspeptin receptor (Kiss1r) signaling on GnRH activity and secretion is fundamental to inducing LH release (Clarkson and Herbison, 2009), crucial for the establishment and maintenance of reproduction (Roux et al., 2003; Seminara et al., 2003; Funes et al., 2003). However, the need for GnRH/LH pulses and surges, rather than prolonged, sustained GnRH activation due to kisspeptin stimulation, which is incompatible with this function, suggests the existence of an “Off” switch. It is known that kisspeptin neurons of the anteroventral periventricular nucleus (AVPV) are important components of the mechanism mediating the positive feedback effects of estrogen on the GnRH/LH surge, while the subpopulation of kisspeptin neurons in the arcuate nucleus (ARH), frequently referred to as KNDy neurons due to their coexpression of kisspeptin, neurokinin B and dynorphin, has been associated with the control of LH pulsatile secretion during the negative feedback phase of estrogen (Oakley et al., 2009; Starrett and Moenter, 2023). While the activatory role of kisspeptin is well-established, the negative regulators of the GnRH system, responsible for ending the kisspeptin response, remain largely understudied. For example, the KNDy neuronal hypothesis can explain the control of kisspeptin neurons themselves but not how GnRH neurons are controlled (Velasco et al., 2023), since dynorphin is not necessary for the termination of the GnRH/LH pulses in rodents (Mostari et al., 2013) and in addition, is lacking in human KNDy neurons (Hrabovszky et al., 2012). This knowledge gap leaves numerous questions unanswered regarding the intricate regulatory network governing the GnRH axis and hence the mechanisms that shape the GnRH surge and pulse.

In a recent review, we postulated the existence of a Kisspeptin-nNOS-GnRH or “KiNG” neuronal network that would function by alternating between neuronal activation and tonic inhibition in order to modulate LH pulse and surge generation (Delli et al., 2021). Similar to kisspeptin, neurons expressing the neuronal NO synthase (nNOS, coded by *Nos1*), which produce the highly diffusible gaseous messenger NO, are estradiol-sensitive and express the estrogen receptor alpha (ERα) (Chachlaki et al., 2017b). nNOS neurons are morphologically and functionally associated with GnRH neuronal cell bodies and dendrites in the medial preoptic area (MePO) and the organum vasculosum of the lamina terminalis (OV); NO acts as an inhibitory signal that rapidly integrates and transmits peripheral signals to the brain, particularly to GnRH neurons (Chachlaki et al., 2017a). The importance of NO signaling in the regulation of GnRH functionality is increasingly apparent (for review see (Chachlaki et al., 2017a), and loss-of-function mutations in *Nos1* have recently been identified in a cohort of patients with congenital hypogonadotropic hypogonadism (Chachlaki et al., 2022), demonstrating the key role of nNOS in reproduction. This hypogonadotropic hypogonadal phenotype has also been reported in a mouse model of genetic Nos1 deficiency (Nos1^-/-^) (Gyurko et al., 2002; Hanchate et al., 2012; Chachlaki et al., 2022), which exhibit both abnormal estrous cyclicity and blunted LH levels in proestrus (Chachlaki et al., 2022), while the intracerebroventricular injection of NOS blockers inhibits GnRH/LH secretion (Rettori et al., 1993), suggesting that nNOS activity and NO release are involved in both positive and negative feedback. In fact, estradiol positively regulates nNOS activity in the OV/MePO by promoting the physical association of the nNOS enzyme with the NMDA receptor (d’Anglemont de Tassigny et al., 2009) and subsequently, phosphorylation of the nNOS enzyme at Ser-1412 (Parkash et al., 2010), an action that requires Kiss1-Kiss1r signaling (Hanchate et al., 2012). Kisspeptin/NO interactions have also been identified as a mechanism shaping GnRH pulsatility by modulating the refractory period of GnRH neuronal activity in response to kisspeptinergic stimuli (Constantin et al., 2021), supporting the so-called KiNG neuronal network hypothesis, i.e. the possibility that a microcircuit of excitatory and inhibitory inputs from kisspeptin and nNOS neurons could orchestrate GnRH neuronal activity and release (Delli et al., 2021).

In this report, we challenge this hypothesis by focusing on the preoptic OV/MePO nNOS population (nNOS^OV/MePO^), surrounding the GnRH cell bodies. Using *in situ* hybridization and various *in vivo* genetic and chemogenetic approaches, we examine the role of the nNOS^OV/MePO^ neurons in the control of pulsatile LH secretion and kisspeptin-induced LH secretion, while also uncovering new mechanistic details of the KiNG neuronal network thanks to the engineering of a new viral tool that for the first time measures *in vivo* NO/cGMP release.

## RESULTS

### Hypothalamic preoptic neurons expressing *Nos1* mRNA differentially express ***Kiss1r* mRNA depending on the estrous cycle phase.**

nNOS enzymatic activity in the OV/MePO, and hence NO levels, have been suggested to be controlled by the classic Kiss1/Kiss1r/AKT pathway (Hanchate et al., 2012), being low during diestrus I (Di) and reaching their maximum during proestrus (Pro) (Parkash et al., 2010). In order to more accurately analyze the anatomical relationship of neurons expressing *Kiss1*, *Kiss1r* and *Nos1* mRNA in the hypothalamic preoptic area, and evaluate plausible differential dynamics across the estrous cycle, we took advantage of the RNAscope technology. Fluorescence in situ hybridization in the OV and MePO of cycling females in diestrus and proestrus revealed the expression of *Kiss1r* mRNA in a fair proportion of *Nos1*-positive cells (Fig. 1 a, b, Suppl. Fig. 2 a). Interestingly, even though the percentage of *Nos1-* or *Kiss1r*-expressing cells remained unaltered (Fig. 1 c, d), the percentage of *Nos1*-expressing cells positive for *Kiss1r* mRNA was significantly increased in proestrus (OV: 45.43% ± 1.80 in Di vs. 55.08% ± 1.322 in Pro, p= 0.04 and MePO: 41.49% ± 4.216 in Di vs. 56.21% ± 2.703 in Pro, p= 0.04) (Fig. 1 b).

**Figure 1.**
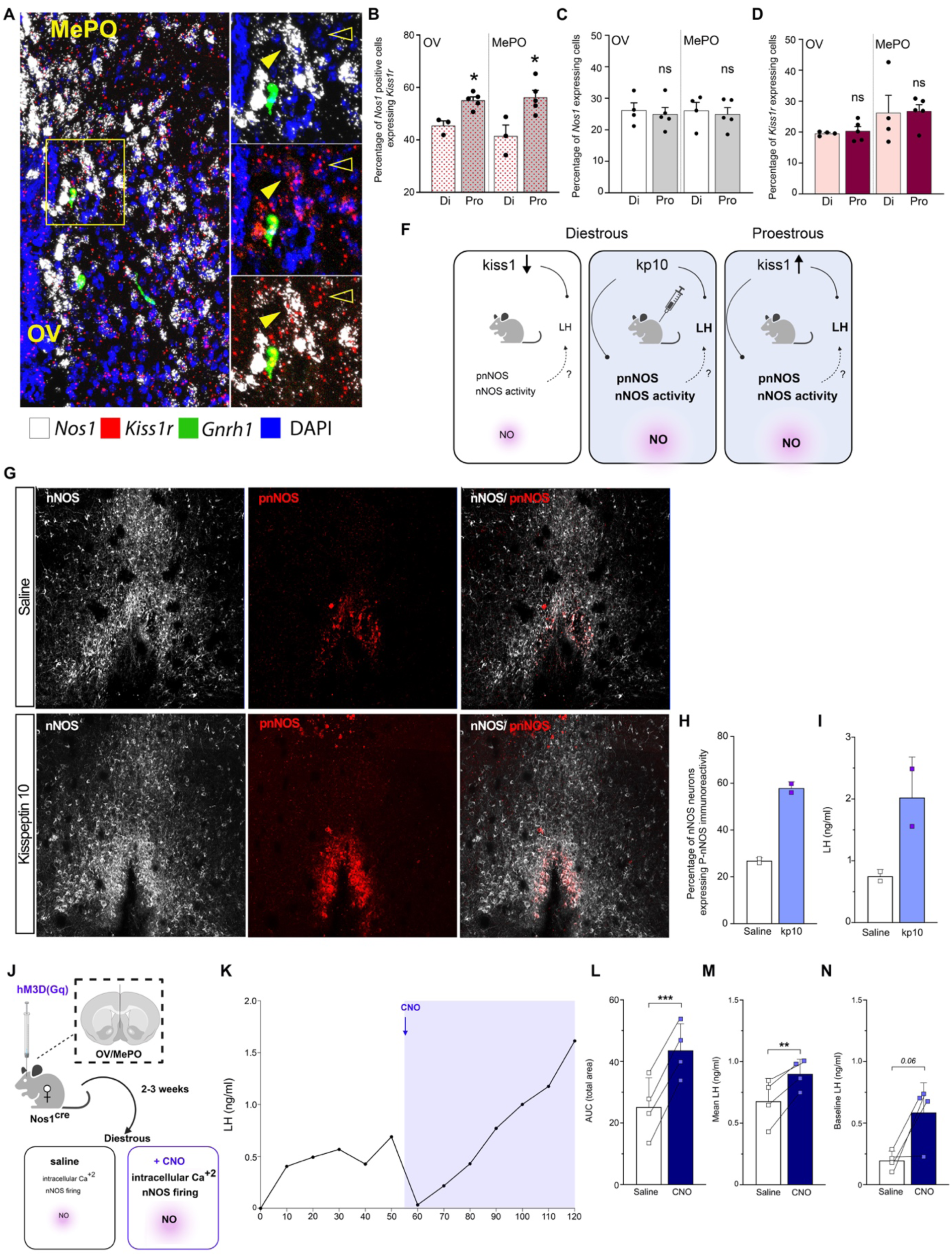
Kisspeptin-nNOS interaction dynamics in the preoptic area of the cycling female mouse. (A-D) Anatomical interaction and differential expression of *Nos1*, *Kiss1* and *Kiss1r* mRNA in the female mouse preoptic area across the estrous cycle. (A) Confocal florescent RNAscope reveals the expression of *Kiss1r* mRNA (red) expression in cells co-expressing variable signals of *Nos1* mRNA (white) in the region of the OV/MePO during proestrous. Magnifications on the right side of the illustrations indicate representative examples of *Nos1* mRNA positive cells co-expressing the *Kiss1r* mRNA (yellow arrowhead), also co-expressed by *Gnrh1*-positive neurons (green) as well as *Nos1*-negative cells (empty arrowhead). (B) Quantification of the percentage of cells positive for the *Nos1* mRNA also being positive for the *Kiss1r* mRNA in Diestrous and Proestrous in the region of the OV and MePO (unpaired t-test; Di, N=3 ; Pro N=5). (C, D) Quantification of the percentage of the cells (over total DAPI) positive for the (C) *Nos1* and (D) *Kiss1r* mRNA in Diestrous and Proestrous in the region of the OV and MePO (unpaired t-test; Di, N=4 ; Pro N=5). (E-H) Kisspeptin effect on nNOS activity in the mouse preoptic area. (E) Graphic illustrating the proposed mechanism of action of physiological kisspeptin fluctuations during diestrous or proestrous, or of the peripheral kisspeptin-10 administration on nNOS activity (F) Immunolabeling for nNOS (white) and P-nNOS (red) in the OV/MePO of female mice upon intraperitoneal administration of saline (top panel) or kisspeptin-10 (bottom panel), (G) quantification of the ratio of P-nNOS ir cells on the total number of nNOS cells and (H) measurement of the circulating LH in mice before (pre) and 30 minutes after (post) receiving a peripheral injection of kisspeptin-10. (I-M) Effect of chemogenetic activation of nNOS neurons in the OV/MePO on LH release pattern. (I) Graphical illustration of the experimental groups. (J) Representative graph of the LH release pattern over the 120 minutes of serial blood sampling. Arrow indicates the time of CNO administration (55^th^ minute). Quantification analysis of the (K) area under the curve, (L) mean and (M) basal LH levels of the Nos1cre animals before (white bar) and after (blue bar) CNO administration. ns: non-significant, * P < 0.05. 3V; third ventricle; AVPV, anteroventral periventricular nucleus; OV, organum vasculosum laminae terminalis; MePO, median preoptic nucleus; Diestrous, Di; Proestous, Pro; kp10, kisspeptin-10.

These changes were specific for the nNOS population of the OV/MePO (nNOS^OV/MePO^). In the hypothalamic anteroventral periventricular nucleus (AVPV), the percentage of *Nos1*-expressing cells co-expressing *Kiss1r* mRNA was found to be unaltered across the estrous cycle (41.85% ± 1.788 in Di vs. 44.,47% ± 2.249 in Pro) (Suppl. Fig. 1 a, c, and Suppl. Fig. 2 b). Even though *Kiss1* and *Nos1* mRNAs were found to be separately expressed in the vast majority of cells in the AVPV (Suppl. Fig. 1 b, g), about 6% of *Kiss1*-positive cells were seen to express low amounts of *Nos1* mRNA at both stages of the estrous cycle (7.795% ± 2.104 in Di vs. 6.061 % ± 1.600 in Pro), with *Kiss1* mRNA showing an almost 50% increase in proestrus (0.1317 % ± 0.01823 in Di, vs. 0.2132% ± 0.01786 in Pro) (Suppl. Fig. 1 f).

### Kisspeptin induces NO-dependent cGMP production in OV/MePO neurons, including GnRH neurons

Previous reports have revealed that the intraperitoneal administration of kisspeptin10 (kp10) promotes an increase in Ser1412-phosphorylation of nNOS in the OV/MePO, accompanied by an increase in LH levels (Hanchate et al., 2012), in keeping with the aforementioned frequent expression of *Kiss1r* in these neurons (Fig.1). Ser1412-phosphorylation is believed to be associated with enhanced nNOS enzymatic activity (Rameau et al., 2007). However, the association of Ser1412-phosphorylation of nNOS with hypothalamic NO production has not been demonstrated *in vivo*. We therefore engineered an AAV9 viral tool expressing the FlincG3 cGMP-dependent biosensor (Fig. 2 a), which we have previously used to accurately measure the levels of NO released by transfected cells *in vitro* (Bhargava et al., 2013; Chachlaki et al., 2022), under the control of the CMV promoter (Fig. 2 b).

**Figure 2.**
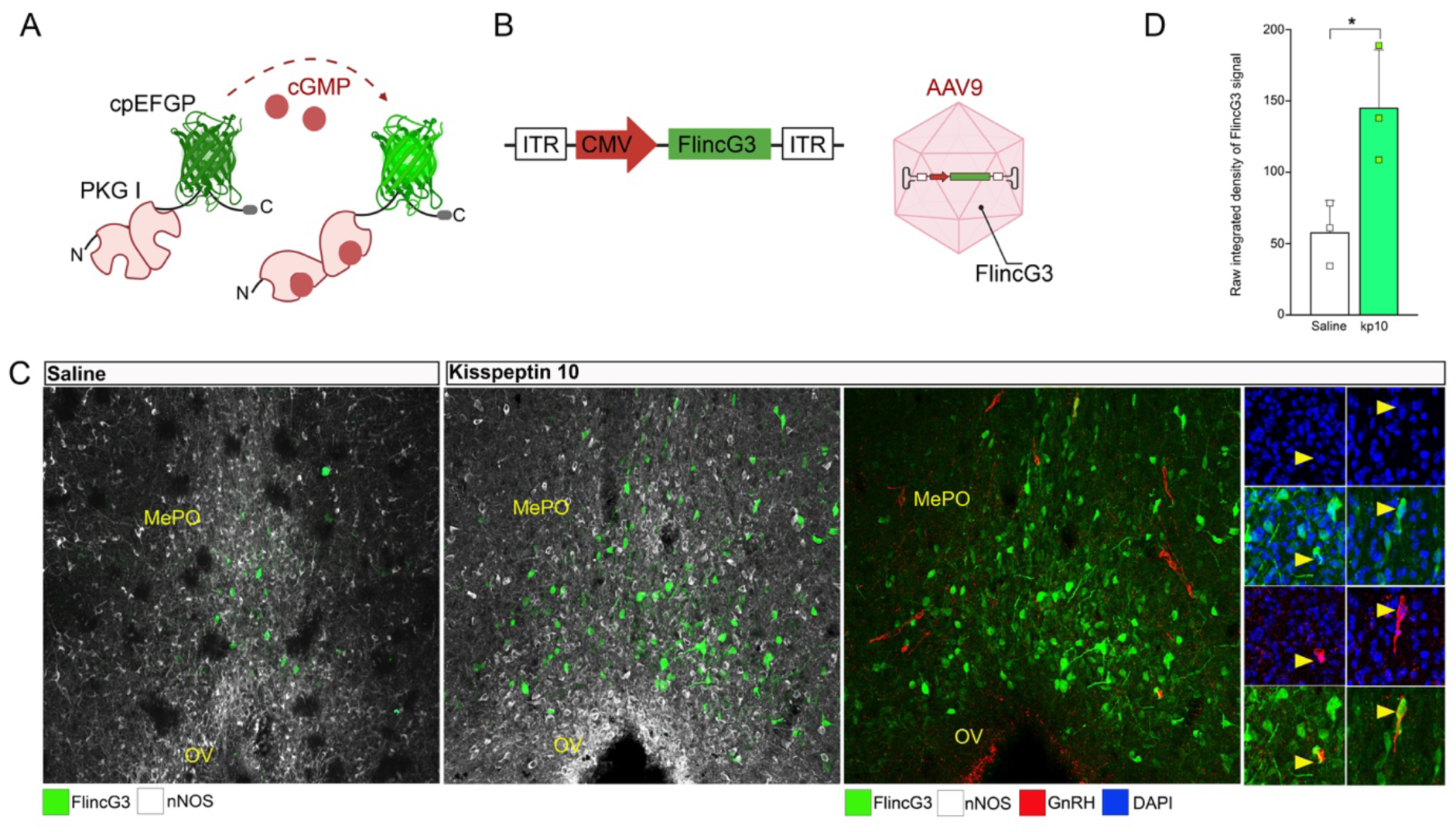
Kisspeptin10 induce the activation of nNOS neurons and the production of cGMP. (A) Illustration of the molecular functioning of the FlincG3 protein and (B) the construction of the viral vector driving its expression. (C) Confocal image of the OV/MePO of wildtype females in diestrous, having received an i.c. injection of AAV9-FlincG3 in the OV/MePO showing the expression of nNOS (white), GnRH (red) and DAPI nuclear staining (blue) along with the endogenous fluorescence of the FlicG3-cpEGFP protein (seen in green) upon an intraperitoneal injection of saline (left panel) or kisspeptin 10 (right panel). Magnifications on the right side of the illustrations indicate representative examples of GnRH positive cells showing FlincG3 activation (yellow arrowhead) upon kisspeptin10 administration. (D) Quantification of the integrated intensity of the endogenous fluorescence of the FlicG3 protein in the OV/MePO, upon application of saline or kisspeptin 10. * P < 0.05. OV, organum vasculosum laminae terminalis; MePO, median preoptic nucleus; kp10, kisspeptin 10.

In order to determine whether cGMP is synthesized in cells different to those expressing the nNOS protein and thus whether NO acts in a paracrine manner to stimulate the synthesis of cGMP, we analyzed FlincG3 fluorescence and the anatomical distribution of cGMP-producing cells in the OV/MePO of cycling females after local stereotactic injections of the FlincG3 construct. Females were perfused during diestrus, when nNOS enzymatic activity and NO release are at their nadir, after a bolus intraperitoneal injection of saline or kp10. Kp10 administration led to an increase in Ser1412-phosphorylation of the nNOS protein in the OV/MePO, as well as circulating LH levels, in agreement with previous reports (Hanchate et al., 2012) (Fig. 1h). Kisspeptin treatment was also accompanied by massive cGMP production in the OV/MePO (Fig. 2 c) as testified by the 200% increase in FlincG3 fluorescence (Fig. 2 d). Interestingly, FlincG3 fluorescence was rarely seen in nNOS-immunoreactive cells (OV: 1.6 % ± 0.5 and MePO: 4.0 % ± 0.7 of nNOS neurons showing FlincG3 signal) but was instead observed in cells surrounding nNOS-expressing neurons, including, strikingly, GnRH neurons (Fig. 2c). Our observations demonstrate for the first time that the phosphorylation of nNOS protein at Ser1412 is coupled with cGMP production *in vivo*, and further support the hypothesis that kisspeptin, by modulating nNOS enzymatic activity, can directly increase local NO/cGMP levels. Altogether, these results suggest the existence of a kisspeptin/NO/GnRH communication network in the OV/MePO.

### Activation of nNOS neurons of the OV/MePO is sufficient to trigger GnRH/LH surge-like release

According to the mathematical model we have previously proposed (Bellefontaine et al., 2014), the potential cumulative activation of nNOS^OV/MePO^ neurons, such as we have shown to occur in the presence of kisspeptin, could result in the build-up of NO to levels capable of influencing neighboring GnRH neurons and synchronizing their activity. In order to assess whether the activation of nNOS^OV/MePO^ neurons is sufficient to induce GnRH/LH release, we stereotaxically delivered a viral vector carrying a designer receptor exclusively activated by designer drugs (DREADD), AAV-hSyn-DIO-hM3D(Gq)-mCherry, into the MePO of Nos1cre (Nos1^cre^, B6.129-Nos1tm1(cre)Mgmj/J) female mice, allowing the selective expression of the activatory DREADD hM3D(Gq) in nNOS^OV/MePO^ neurons (Fig. 1 i). Tail blood was collected every 10 min for a total duration of 120 minutes during diestrus, when nNOS^OV/MePO^ activity is minimum. During the test, mice received an intraperitoneal injection of saline 5 minutes before the pulsatile assessment of LH release (min -5) and a CNO injection to activate the DREADD 1 hour later, at minute 55 (Fig. 1 j). As previously reported for intact female mice during diestrus, basal LH (0.2 ng/ml ± 0.04) and total LH, as assessed by the area under the curve (AUC), were low, with mice displaying one pulse of LH over the first 60 min of blood sampling (Fig. 1 j-m). Interestingly, the selective activation of nNOS^OV/MePO^ neurons upon CNO administration led to a significant increase in basal LH (0.6 ng/ml ± 0.1) (Fig. 1 m), as well as of total LH (Fig. 1 k), approaching values reported during the LH surge. As expected, given that during diestrus NO release in the OV/MePO is very low, the selective inhibition of nNOS^OV/MePO^ neurons in Nos1^cre^ mice stereotaxically injected with AAV-hSyn-DIO-hM4D(Gi)-mCherry carrying an inhibitory DREADD hM4(Gi) had no effect on mean LH, total LH or the overall LH secretion pattern in either male or female mice during diestrus (Suppl. Fig. 3). Overall, these data suggest the intriguing possibility that the kisspeptin-induced activation of nNOS^OV/MePO^ neurons may be sufficient to promote the GnRH/LH surge.

### Neuronal NO plays a role in the kisspeptin-mediated regulation of GnRH neuronal activity

Previous studies have suggested that NO modulates the excitability of GnRH neurons via the sGC-cGMP signaling cascade (Clasadonte et al., 2008). Our results reveal that nNOS^OV/MePO^ neurons are the source of massive NO-induced cGMP production in the vicinity of GnRH neurons, including by GnRH neurons themselves, that could lead to physiological GnRH/LH secretion. To explore whether NO deficiency could impact GnRH neuronal activity, we performed electrophysiological recordings of GnRH neurons in the hypothalamus of diestrous females upon pharmacological or genetic manipulation of the nNOS/NO signaling cascade, using a Nos1-deficient transgenic mouse line (Nos1^−/−^). In agreement with what has been previously reported in infantile mice (Chachlaki et al., 2022), spontaneous firing in GnRH neurons was markedly increased in adult female *Gnrh*::*Gfp*; *Nos1*^-/-^ mice, compared to their *Gnrh*::*Gfp*; *Nos1*^+/+^ littermates (Suppl. Fig. 4). Interestingly, this sustained exacerbated GnRH neuron firing (from minipuberty until adulthood) is associated with significantly reduced total and mean LH levels in both female and male Nos1^−/−^ mice (Suppl. Fig 5), in agreement with the hypogonadotropic hypogonadal phenotype associated with nNOS deficiency (Chachlaki et al., 2022; Delli et al., 2023).

Our results illustrated in Figure 2 suggest that kisspeptin directly acts on nNOS^OV/MePO^ neurons to modulate NO/cGMP release across the estrous cycle, and possibly reveal the existence of a local kisspeptin/nNOS network controlling GnRH neuronal activity and pulsatile release pattern. Bath application of kp10 was able to stimulate GnRH firing in hypothalamic slices from both *Gnrh*::*Gfp*; *Nos1*^+/+^ and *Gnrh*::*Gfp*; *Nos1*^-/-^ mice (Fig. 3 a, b, e). However, genetic nNOS deficiency resulted in exacerbated kp10-induced GnRH neuronal firing (Fig. 3 d, e). Similarly, the pharmacological blockade of NO production via the *ex vivo* application of the NOS inhibitor L-NAME resulted in an enhanced kp10-induced increase in GnRH neuron firing, reaching levels comparable to those seen in *Nos1*^-/-^ mice (Fig. 3 a-b, d, e). Intriguingly, the e*x vivo* bath application of an NO donor (DEA/NO) to the OV/MePO abolished this exacerbated kp10-induced response of GnRH neurons in *Gnrh*::*Gfp*; *Nos1*^-/-^ mice, reducing the firing rate to those seen in *Gnrh*::*Gfp*; *Nos1*^+/+^ mice upon kp10 application (Fig 3 e). Hence, our results further support a role for NO as a negative regulator of GnRH neuronal activity, and that the kisspeptin-induced electrical response of GnRH neurons is fine-tuned by NO production from nNOS^OV/MePO^ neurons. These results suggest that the action of kisspeptin on GnRH neuronal activity is finely tuned by the locally induced release of nitric oxide.

**Figure 3.**
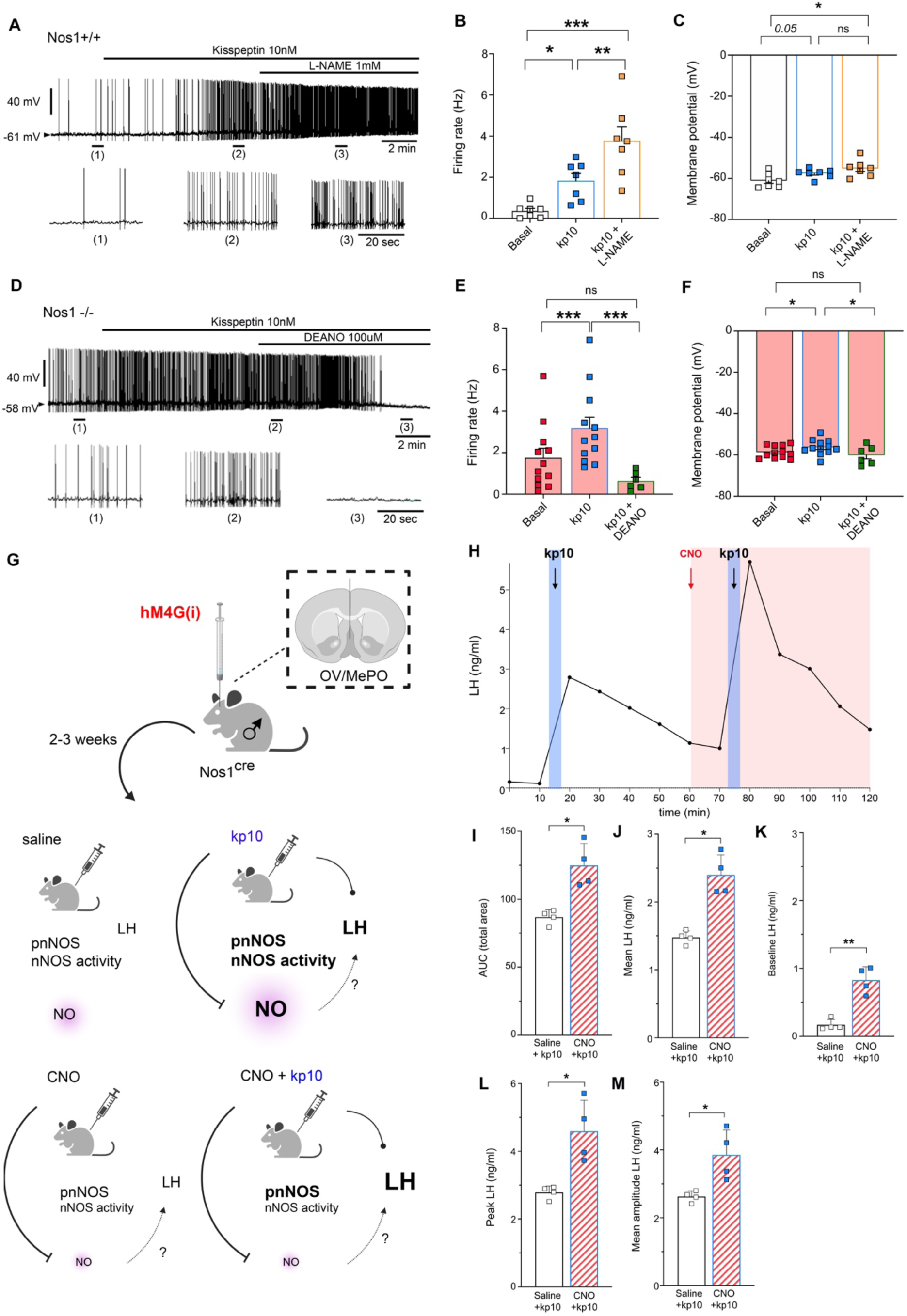
nNOS ^OV/MePO^ NO signalling controls GnRH neuronal activity and release in kisspeptin-excited GnRH neurons. (A) Whole-cell current-clamp recording of a GnRH neuron from a *Nos1*^+/+^ animal in the presence of bath application of kisspeptin (10 nM) and the NOS inhibitor (L-NAME 1 mM). Bottom traces show expanded timescales of the recording at the indicated time point 1,2 and 3. (B-C) Quantification of (B) spontaneous firing frequency and (C) membrane potential in GnRH neurons from *Gnrh::Gfp; Nos1*^+/+^ under basal conditions (white), in the presence of kisspeptin10 (blue) and the co-application of L-NAME (yellow) (One way repeated measures ANOVA followed by Tuckey’s post hoc test, n=7 cells, N= 6 mice). (D) Whole-cell current clamp recording of a GnRH neuron from a *Nos1*^-/-^ animal in the presence of bath application of kisspeptin (10nM) and the NO donor (DEA/NO 100 uM). (E) Average firing rate and (F) membrane potential of GnRH neurons recorded from *Gnrh::Gfp; Nos1*^-/-^ mice (red) under basal conditions (red), in the presence of kisspeptin10 (blue) and the co-application of DEANO (green) (Mixed-effects analysis followed by Tuckey’s post hoc test; Basal and Kisspeptin, n=12 cells, N=6 mice; DEA/NO, n=6 cells, N=4 mice). (G) Graphical illustration of the experimental groups of intact Nos1cre male mice injected into the MePO with the AAV-hSyn-DIO-hM4(Gi)-mCherry and intraperitoneally with kp10 (1nmole, 15^th^ and 75^th^ minute). (H) Representative LH pulse profile and subsequent analysis. (I) Area Under the Curve (AUC) analysis, (J) Mean total LH, (K) Mean baseline LH levels, (L) Peak LH response, and (M) Mean amplitude of LH responses before (white) or after (purple) CNO administration (60^th^ minute). kp10, kisspeptin 10. N=4, ns: non-significant, Values indicate means ± SEM * P < 0.05, ***P<0.001.

### nNOS neurons of the OV/MePO are integral modulators of the kisspeptin-mediated GnRH/LH release

Since, according to our results, kisspeptin promotes NO/cGMP production that in turn is required for the precise regulation of the kisspeptin-induced GnRH neuronal activity, we focused our investigations on whether nNOS^OV/MePO^ neuronal activity could itself be involved in the fine-tuning of kisspeptin-mediated GnRH/LH release. To this end, we first selectively injected the DREADD hM4(Gi) virus into the MePO of Nos1^cre^ male mice, the activation of which upon CNO administration had no effect on GnRH/LH release in either female or male mice (Suppl. Fig. 3). Kp10 was then intraperitoneally administered before and after CNO administration (Fig. 3 g, h). As expected, the application of kp10 under basal nNOS^OV/MePO^ neuronal activity (i.e., before CNO) induced a rapid increase in LH release (peak LH= 2.79 ng/ml ± 0.09) (Fig. 3 h). Intriguingly, the inhibition of the nNOS^OV/MePO^ population resulted in an exacerbated kp10-induced LH response upon the administration of CNO (peak LH= 4,59 ng/ml ± 0.46) (Fig. 3 h-m). Total (Fig. 3 i), mean (Fig. 3 j) and basal LH levels (Fig. 3 k) were all increased in the presence of kisspeptin upon CNO administration. These results together with our electrophysiological data, strengthen the importance of nNOS^OV/MePO^ neuronal activity in the fine-tuning of kisspeptin-induced GnRH activity and subsequent LH secretion.

### Kisspeptin shapes GnRH/LH release through a dual activation-inhibition mechanism involving NO

In order to further explore the physiological significance of kisspeptin/NO interactions for the GnRH secretion profile, we assessed whether *in vivo* pharmacological manipulation of NO/cGMP signaling could modulate the kisspeptin-induced LH response. To this end, intact WT male mice were intraperitoneally administered saline, L-NAME or sildenafil – a selective enhancer of the NO-induced production of cGMP – 30 minutes before blood sampling, and were then subjected to two kp10 administrations at an interval of one hour (Fig. 4 a). As expected, kp10 administration induced a rapid, sharp increase in LH secretion in saline-treated animals. The administration of L-NAME did not impact the ability of kp10 to efficiently stimulate GnRH/LH secretion, since, within 5 minutes of kp10 administration, both groups had identical LH levels (Fig. 4 f). Yet, in the absence of NO, the LH profile during the resolution phase was significantly altered. In agreement with what was seen upon pharmacogenetic inhibition of nNOS^OV/MePO^ neurons, the administration of L-NAME induced an overshooting of the kisspeptin-induced LH response, which continued to rise even when LH values in saline-treated animals were rapidly decreasing (Fig. 4 e), supporting the view that kisspeptin-induced NO release is necessary to end kisspeptin-induced GnRH/LH secretion. Interestingly, the administration of sildenafil, by prolonging the activity of the NO/cGMP cascade, led to a faster clearing of the kisspeptin effect, with LH levels eventually dropping below those in saline-treated animals (Fig. 4 g). A second administration of kp10, at the same dose as the initial one, 60 minutes later, resulted in a potentiated LH response to kisspeptin in animals treated with saline (Fig. 4 f). Animals treated with L-NAME displayed an equally exacerbated LH response to the second kp10 administration, similar to what was seen during the first kp10 administration (Fig. 7 e-f). Intriguingly, sildenafil treatment, by promoting additional NO production and further accumulation of cGMP (due to the inhibition of its breakdown), prevented the potentiation of LH release during the second administration of kp10 compared to the first one, leading to an attenuated peak value of LH levels in comparison to the saline-treated animals (Fig. 7 e-f). Of note, sildenafil treatment resulted in a faster decline in LH levels, with lower nadir levels at the end of the experiment (Fig. 7 g). Overall, our data support a role for NO/cGMP signaling in both amplification and resolution phases of the kisspeptin-induced LH response.

**Figure 4.**
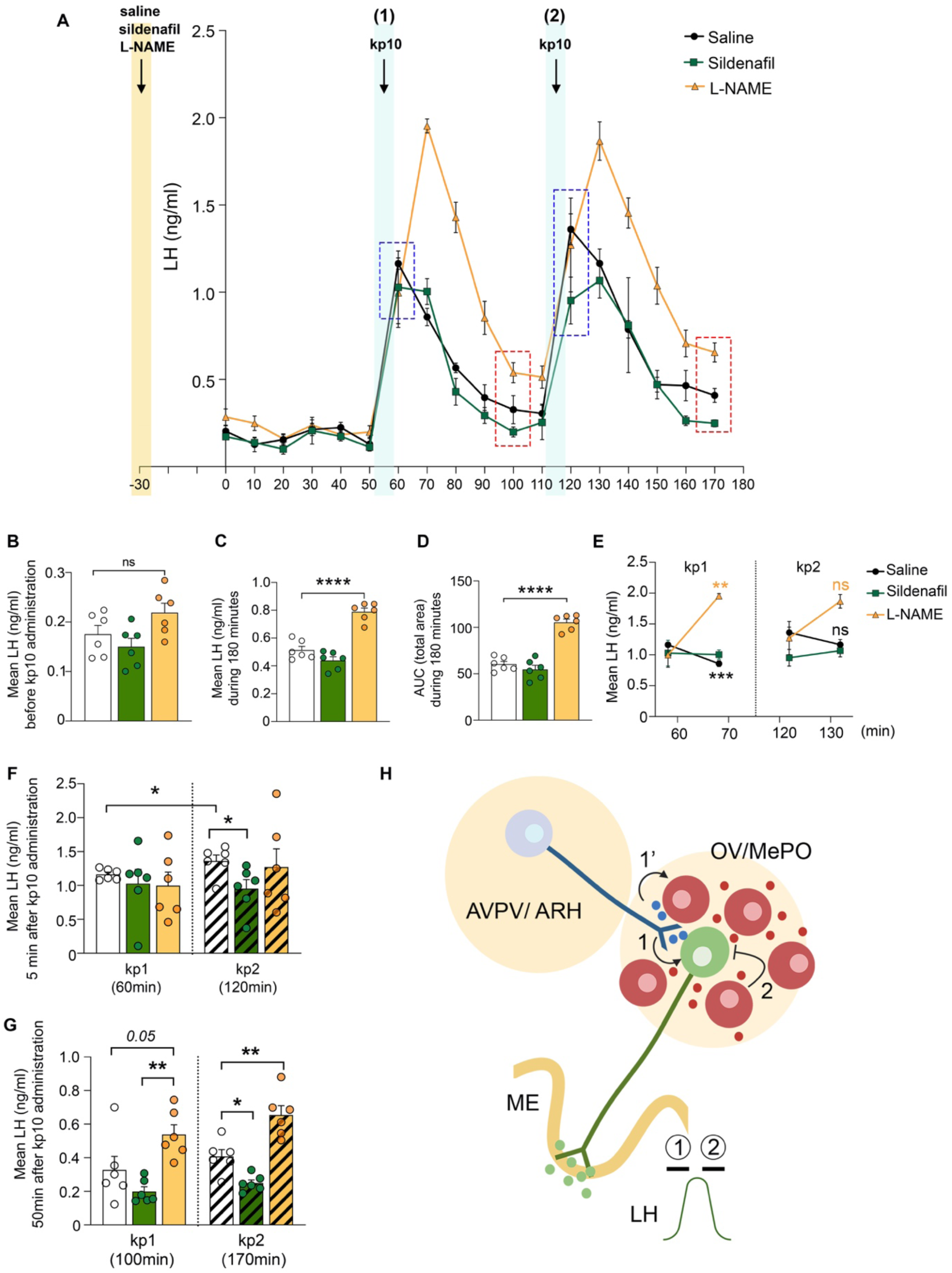
Effect of NO/cGMP signalling in the kisspeptin – induced LH response. (A) Mean LH pulse profiles of 6 intact WT male mice after the administration of saline (black), L-NAME (yellow), or Sildenafil (green), 30 minutes before the beginning of the blood collection (yellow trace). Kisspeptin-10 (kp10; 1nMol) was administered at min 55 and 115 (blue traces marking kp10 administration). (B) Mean baseline LH levels before administration of kisspeptin10. (C) Mean total LH and (D) Area Under the Curve (AUC) analysis for the total duration of 170 min (E) Comparison of mean LH levels 5 minutes vs10 minutes after the first (kp1; minute 60 vs minute 70) and second (kp2; minute 120 vs minute 130) kisspeptin 10 administration. (F) Mean LH levels 5 minutes upon first (kp1; minute 60) and second (kp2; minute 120) kisspeptin 10 administration. (G) Mean LH response at washout upon the first (kp1; minute 100) and the second (kp2; minute 170) kisspeptin 10 administration. (H) Graphical illustration explaining the possible mechanism behind the observed kisspeptin/NO interactions on the fine-tuning of GnRH/LH release. (1) Activation of the kisspeptin neurons of the AVPV and/or ARH (blue) drives GnRH/LH release by (1) acting on the GnRH neurons, while in parallel (1’) stimulates NO release from nNOS^OV/MePO^ neurons (red). (2) NO production in the vicinity of the GnRH neurons (green) will eventually inhibit the GnRH firing through the production of cGMP, ending thus the kisspeptin-induced response. AVPV, anteroventral periventricular nucleus; ARH, arcuate nucleus; OV, organum vasculosum laminae terminalis; MePO, median preoptic nucleus; kp10, kisspeptin 10. N=6, ns: non-significant, Values indicate means ± SEM * P < 0.05, ***P<0.001.

## DISCUSSION

The identification and characterization of both the positive and the negative regulators of the GnRH neural network is crucial for understanding the intricate balance of stimulatory and inhibitory signals that govern reproductive function. Kisspeptin is the most potent stimulator of GnRH activity, while neuronal NO acts on GnRH neurons as an inhibitory regulator (for review see Chachlaki et al., 2017a; Starrett and Moenter, 2023).

Here we report that the nNOS^OV/MePO^ neurons express the *Kiss1r* gene, and that the proportion of *Kiss1r*-expressing nNOS neurons varies across the estrous cycle, with the highest expression occurring in proestrus. Using newly-engineered state-of-the-art sensor probes, we have demonstrated that kisspeptin, by enhancing nNOS enzymatic activity, stimulates the production of NO from nNOS^OV/MePO^ neurons *in situ. In vivo* experiments involving the peripheral administration of kisspeptin and the chemogenetic manipulation of nNOS^OV/MePO^ neurons reveal that kisspeptin fine-tunes LH release by modulating nNOS neuronal activity. Additionally, pharmacological approaches used to either suppress or exacerbate kisspeptin-induced NO release indicate that kisspeptin uses the highly diffusible NO to cap the amount of LH released and the shutdown of the triggered LH surge. Intriguingly, we observed a similar phenomenon at the cellular level, where kisspeptin was seen to use NO to contain the firing activity of the GnRH neurons and potentially terminate its own effects.

This tight relationship of nNOS and kisspeptin neurons and their ability to modulate GnRH activity in opposite ways supports the KiNG (Kisspeptin/ nNOS/ GnRH) neuronal network hypothesis, which indicates the existence of a kisspeptin/ nNOS microcircuit of excitatory and inhibitory inputs coordinating GnRH/LH release. Indeed, deleterious mutations in the genes encoding Kiss1 or Kiss1r (de Roux et al., 2003; Topaloglu et al., 2012), and more recently Nos1 (Chachlaki et al., 2022), have all been found in patients with congenital hypogonadotropic hypogonadism, i.e., GnRH deficiency, supporting the importance of both kisspeptin and NO signaling in the control of fertility.

The expression of *Kiss1r* by non-GnRH neurons has been a matter of debate, possibly due to the low sensitivity of the techniques previously used to detect *Kiss1r* in these neurons (Herbison et al., 2010). However, the observation that the reinstatement of *Kiss1r* expression specifically in GnRH neurons of *Kiss1r* ^-/-^ mice is not sufficient to restore the LH response upon kisspeptin administration (Leon et al., 2016) supports the involvement of other, non-GnRH, Kiss1r-expressing cells in the shaping of the kisspeptin-induced LH release. This is in agreement with our data showing that peripheral kisspeptin administration induces a 100% increase in the phosphorylation levels of the nNOS protein at the Ser1412 activation site and actually increases the local production of NO and of cGMP in neurons surrounding the nNOS^MePO^ neurons. This aligns with the previously reported role of Kiss1/Kiss1r in the mechanism driving the estradiol-promoted enzymatic activation of nNOS occurring on the day of proestrus in the OV/MePO, correlating increased nNOS activity with the GnRH/LH surge (Parkash et al., 2010; Hanchate et al., 2012). The ability of nNOS neurons to directly respond to kisspeptinergic signals in an estrogen-dependent manner supports the KiNG neuronal network hypothesis, suggesting that GnRH release could be ultimately determined by the equilibrium resulting from the interplay of NO and kisspeptin signaling responses across the estrous cycle. High estrogen levels during proestrus, by increasing the AVPV kisspeptinergic inputs to nNOS^OV/MePO^ neurons and thus promoting their enzymatic activation, could have a stimulatory effect on both nNOS and kisspeptin neurons, subsequently enabling the GnRH/LH surge. Indeed, the knockdown of ERα in AVPV kisspeptin neurons blunts the LH surge, maintaining LH pulsatile secretion (Wang et al., 2019; Clarkson et al., 2023), in agreement with the lack of effect of nNOS^OV/MePO^ inhibition on the GnRH/LH profile during diestrus, and previous studies suggesting that nNOS activity is dispensable for continuous/basal release but is required for stimulated GnRH/LH release (Bonavera et al., 1993). In fact, according to the hypothesis we have previously presented, based on realistic mathematical modelling of the size and distribution of nNOS neurons in the OV/MePO (Bellefontaine et al., 2014), when the enzymatic activity of nNOS^OV/MePO^ neurons is massively increased, NO concentration could potentially rise to levels capable of influencing neighboring GnRH neurons and synchronizing their activity. Yet, even though the stimulatory role of nNOS Ser1412 phosphorylation has been coupled with rapid NO production *in vitro* (Rameau et al., 2007), such a mechanism has never been demonstrated in the hypothalamus *in vivo*. By generating a powerful new tool that accurately measures NO/cGMP concentration *in vivo*, our study is the first to demonstrate that the kisspeptin-induced nNOS phosphorylation is actually associated with massive NO/cGMP production in the OV/MePO of both sexes. This kisspeptin-induced NO production is additionally coupled with an increase in LH levels, supporting the view that the highly diffusible NO is the perfect candidate to synchronize the widely dispersed GnRH neuronal population, thus enabling the GnRH/LH surge. In support to this hypothesis, our results reveal that the pharmacogenetic activation of nNOS^OV/MePO^ neurons during diestrus, i.e., when nNOS activity is at its nadir, is sufficient to provoke a proestrous-like LH surge, in agreement with previous studies showing that central NMDA administration can elicit LH release in Kiss1^-/-^ and Kiss1r^-/-^ mice, acting at least partially through nNOS neurons (d’Anglemont de Tassigny et al., 2010). Indeed, our results, by demonstrating a paracrine action of NO and most importantly, the production of cGMP by GnRH neurons themselves, further reinforce the idea that NO could be a part of the mechanism modulating GnRH/LH release by acting downstream of kisspeptin, directly on GnRH neurons.

Previous studies had already provided evidence that NO could modulate GnRH neuronal activity via a mechanism involving soluble guanylate cyclase (sGC)-induced cGMP production (Clasadonte et al., 2008; Chachlaki et al., 2022). According to our electrophysiological recordings of GnRH neurons, alterations of NO release in the vicinity of GnRH neurons can impact GnRH neuronal activity not only directly but also indirectly, by affecting the ability of peripheral kisspeptin to modulate the GnRH neuronal activity pattern. In fact, genetic or pharmacologically-induced NO deficiency was seen in our study to amplify the kisspeptin-induced GnRH firing activity, while excess NO resulted in a drop in kisspeptin-induced GnRH neuronal firing frequency, suggesting that nNOS^OV/MePO^ neuronal activity and NO homeostasis are necessary components of the kisspeptin-regulated mechanism responsible for the fine-tuning of the GnRH firing pattern and subsequent GnRH release. A recent study in fact indicates that kisspeptin/NO interactions provide a mechanism for the modulation of the refractory period of GnRH neuronal activity (Constantin et al., 2021). One could hypothesize that after activating GnRH neurons, kisspeptin, by promoting NO release, activates a brake, ensuring that the GnRH system doesn’t switch to a sustained, surge mode of GnRH/LH release. Indeed, the pharmacogenetic inhibition of nNOS^OV/MePO^ activity or the pharmacological inhibition of NO by L-NAME, by suppressing the inhibitory action of NO, led to an exacerbated kisspeptin-stimulated LH response. Importantly, this NO blockade did not alter the ability of kisspeptin to directly promote the release of GnRH, but rather promoted a prolonged rise of LH levels, suggesting an inefficient arrest of GnRH stimulation, similar to what has been previously shown upon the ablation of KNDy neurons in the rat brain (Helena et al., 2015). Actually, previous studies have already reported the presence of a subset of KNDy neuron afferents displaying oscillatory activity and projecting to the preoptic area (Yeo and Herbison, 2011), where their projections are apposed to both GnRH and nNOS neurons (Bouret et al., 2004; Caron et al., 2012), raising the possibility that ARH neurons could also be involved in the indirect control of the nNOS^OV/MePO^ population. In accordance with the ability of exogenous NO to induce a decrease in kisspeptin-induced GnRH firing, the administration of sildenafil, by triggering a gradual accumulation of excess cGMP around GnRH neuronal somata, abrogates the ability of kisspeptin to elicit a progressive increase in the magnitude of the LH response, a phenomenon previously reported in rats (Tovar et al., 2006), eventually resulting in lower basal LH levels. More specifically, under these conditions, the first kisspeptin10 (kp10) administration induces the activation of GnRH neurons and hence the secretion of GnRH from its nerve terminals, and in parallel, activates nNOS^OV/MePO^ neurons, resulting in NO-induced cGMP release, GnRH inhibition and thus the termination of kisspeptin-induced LH secretion. In the presence of sildenafil, cGMP levels remain constantly high, prolonging its effect. A second kp10 administration would thus have led not only to a reactivation of GnRH neurons, but also to the further accumulation of cGMP due to another round of nNOS^OV/MePO^ activation. The excess cGMP levels thus lead to the hyper-inhibition of GnRH neurons, negatively impacting LH secretion. Overall, these data strongly suggest that the fine-tuning of nNOS activity, and hence NO levels, is a crucial part of the mechanism shaping the GnRH/LH release pattern over the estrous cycle, with low levels of NO being indispensable for the maintenance of GnRH/LH pulsatile release, while the activation of nNOS^OV/MePO^ neurons promotes the LH surge. Our results further support the view that the nNOS-kisspeptin communication network involves the traditional Kiss1r/nNOS/cGMP pathway.

In summary, our study provides evidence supporting the involvement of the KiNG neuronal network in shaping the GnRH/LH release pattern. According to this model, during the estrogen negative feedback phase in diestrus, the activation of KNDy neurons results in kisspeptin release, stimulating GnRH secretion. In parallel, nNOS^OV/MePO^ neurons directly sense kisspeptin signals in a tightly regulated manner due to the differential expression of the *kiss1r*, possibly ensuring the tight regulation of nNOS enzymatic activity in nNOS^OV/MePO^ neurons by kisspeptin. This fine-tuning of nNOS enzymatic activity by kisspeptin allows, during the negative feedback phase, the production of low levels of NO/cGMP in the vicinity of GnRH neurons, which are indispensable for physiological GnRH neuronal activity, in part by switching off kisspeptin-induced GnRH/LH production and allowing the GnRH system to return to baseline. During the positive feedback page, the activation of AVPV kisspeptin neurons would result in kisspeptin being released around the nNOS^OV/MePO^ neuronal population that, being more sensible to the kisspeptin signal now due to increased *kiss1r* expression, becomes maximally activated, allowing massive NO/cGMP release. The direct actions of kisspeptin on GnRH production/release and its indirect actions through NO could be thus two sides of the same coin, acting as the ON and OFF signals to fine-tune GnRH neuronal activity and instigate the LH surge. Aside from their potential physiological relevance, our current results further contribute to the hypothesis that NO could be the long-missing key to the synchronization of GnRH neurons.

## MATERIALS AND METHODS

### ANIMALS

All C57Bl/6J mice were housed under specific pathogen-free conditions in a temperature-controlled room (21-22°C) with a 12h light/dark cycle and ad libitum access to food and water. Experiments were performed on male and female C57BL/6J mice (Charles River Laboratories), Nos1cre mice (B6.129-Nos1tm1(cre)Mgmj/J), Nos1-deficient (B6.129S4-Nos1tm1Plh/J) mice and Gnrh::Gfp mice (a generous gift of Dr. Daniel J. Spergel, Section of Endocrinology, Department of Medicine, University of Chicago, IL) (97). Nos1-/-; Gnrh::Gfp mice were generated in our animal facility by crossing Nos1-/+ mice with Gnrh::Gfp mice. Animal studies were approved by the Institutional Ethics Committees for the Care and Use of Experimental Animals of the Universities of Lille; all experiments were performed in accordance with the guidelines for animal use specified by the European Union Council Directive of September 22, 2010 (2010/63/EU) and were approved by the French Department of Research (APAFIS#2617-2015110517317420v5).

### PHARMACOLOGICAL AGENTS

Synthetic kisspeptin-10 [rodent metastin (45–54) amide; YY-10-NH2] was purchased from GeneCust. The NOS inhibitor N-G-nitro-l-arginine methyl ester, HCl (l-NAME), and DMSO (317275) were purchased from Calbiochem. Sildenafil citrate was purchased from Sigma-Aldrich. The diethylamine (DEA)/NO (2-(N, N-diethylamino)-diazenolate-2-oxide.diethyl-ammonium salt) was from Alexis (San Diego, CA).

### STEREOTAXIC INJECTIONS

The AAV9 hM4D DREADD (AAV-hSyn-DIO-hM4D(Gi)-mCherry) (titer = 4.5 × 10^12^ vg/mL; Addgene; 44362-AAV9) and the AAV9 hM3D DREADD (AAV-hSyn-DIO-hM4D(Gq)-mCherry) (titer = 4.5 × 10^12^ vg/mL; Addgene; 44361-AAV9) was stereotaxically injected into the OV/MEPO of Nos1-cre female and male mice. The AAV9 FlincG3 (AAV-CMVFlincG3) (titer = 10 x 1012 vg/mL; Genecust) was stereotaxically injected into the OV/MEPO of wild-type female mice. Animals were kept under general anesthesia (induction 4% in air 2 L/min, then 1.5% in air 0.3 L/min), after local injection of lidocaine (30 microliters of a 0.5% solution, s.c.) and preemptive Meloxicam treatment (5 mg/kg). Using a 2 μL Hamilton Neuros syringe (Hamilton; catalog # 65459-01), 300 nL of a solution containing virus was delivered at a rate of 80 nL per minute after a 5-min pre-injection period, and followed by a 10-min post-injection period. Mice were left to recover from brain surgery for three weeks, which also allows sufficient transduction of viral vector. To target the OV/MEPO, following coordinates were used: anterior-posterior (A-P) = + 1.3 mm; medial-lateral (M-L) = 0 mm; dorsal-ventral (D-V) = - 4.8 mm. To chemogenetically manipulate nNOS neurons, mice received an intraperitoneal (ip) injection of 100 μL CNO (Tocris; 6329) at a dose of 1 mg/kg diluted in sterile saline.

### FLUORESCENCE IN SITU HYBRIDIZATION

FISH was performed on frozen brain sections of 3 months old female mice with the RNAscope Multiplex Fluorescent Kit v2 (323100, Advanced Cell Diagnostics). Estral cyclicity was assessed during 2 weeks by daily vaginal smears. Animals were then sacrificed by decapitation in the afternoon of the first day of diestrus or proestrus. Brains were dissected and included in OCT medium (Tissue-Tek), frozen on dry ice and stored at −80°C until sectioning, while the edematous swelling of the uterus was visually checked to confirm the estral stage. Tissues were cryosectioned (Leica cryostat) coronally at 16 μm on Superfrost Plus slides and stored at −80°C. Sections were processed according to the manufacturer’s protocol, using specific probes to detect Nos1 (437651-C2, NM_008712.2, target region 2-1097), Kiss1r (408001, NM_053244.5, target region 21 – 1599), Kiss1 (476291, XM_006529679.2, target region 121-1376) and Gnrh1 (476281-C3, XM_006518564.3, target region 81-914) mRNAs. Hybridization with a probe against the Bacillus subtilis dihydrodipicolinate reductase (DapB) gene (320878) was used as a negative control.

RNAscope preparations were analyzed on the LSM 710 Zeiss confocal microscope. Excitation wavelengths of 405/443nm, 488/518 nm, 561/699 and 633/699 were selected to image the different channels. All images were taken with the objective EC Plan-Neofluar M27 (thread type) using the 40X oil objective with a numerical aperture of 0.50, and a zoom of 1.4. Z-stack images were taken every xx μm, for a total of xx μm per sections. All the images are presented as maximal intensity projections (MIP) of three-dimensional volumes along the optical axis. Images to be used for figures were pseudocolored and adjusted for brightness and contrast using Photoshop (Adobe Systems, San Jose, CA).

#### Cell counting

The nomenclature of brain regions used in this article and the definition of the region of interest (ROI) was based on the ones described in the Allen brain atlas (Lein et al., 2007). The cell counts were undertaken unilaterally on maximal intensity projections (MIP) images by counting the number of single-labeled Nos1 positive neurons and double-labeled Nos1/Kiss1r neurons in the region of the OV/MePO (represented by plate 50 of the Allen Mouse Brain Atlas), AVPV (plate 53) and ARH (plate 71). This quantification is represented as percentage of Nos1 neurons co-expressing kisspeptin receptor, and the values from each mouse were averaged to determine mean counts and SEM for each age group. To quantify Kiss1 mRNA content, the integrated density analysis was carried out using ImageJ analysis software. Each image was binarized, so as to isolate labeled neurons from background, as well as compensate for differences in fluorescence intensity. The integrated intensity was then calculated for each image, this reflecting the total number of pixels in the skeletonized image.

### IMMUNOHISTOCHEMISTRY

#### Tissue Preparation

Mice were anesthetized with a lethal dose of pentobarbital (DOLÉTHAL®, 5mg/kg) and locally treated with Lidocain (30 μl of a 0.5% solution). Animals were perfused thoroughly with cold saline, until exsanguination, then perfused with cold fixative solution [4% paraformaldehyde (PFA), in 0.1M PB, pH 7.4]. The brain was extracted and post-fixed with the same fixative solution for 2h at 4°C, then cryoprotected in PB 0.1M 20% sucrose for 24h at 4°C. Afterwards, the brain was embedded in OCT medium (Tissue-Tek), frozen on dry ice, and stored at −80°C until sectioning. Tissues were cryosectioned (Leica cryostat) coronally at 35 μm (free-floating sections).

#### Antibodies

The goat polyclonal anti-nNOS antibody (1:1000; Cat# OSN00004G, Lot#VF3001511) and the rabbit polyclonal (1:500; Cat# PA1-032, Lot# UB274796) were purchased from ThermoScientific. The rat polyclonal anti-GnRH (1:2500; Cat# EH1044) was a generous gift from Professor Hrabovszky (Laboratory of Reproductive Neurobiology, Institute of Experimental Medicine, Budapest, Hungary). The Alexa Fluor 647-conjugated donkey anti-goat secondary antibody (1:500; Cat# A21447, Lot# 2465096) used for nNOS immunolabeling, and 568-conjugated donkey anti-rat secondary antibody (1:500; Cat#?, Lot#) used for GnRH were purchased from Invitrogen.

#### Immunolabeling

Sections were washed 3 times for 5 minutes each in PB 0.1M and then incubated in blocking solution (5% NDS, 0.3% Triton X-100 in PB 0.1M) for 1 hour at room temperature. Sections were incubated for 48-72 hours at 4°C with the primary antibodies diluted in blocking solution, then rinsed 3 times for 5 minutes each in PB 0.1M. The sections were incubated with secondary antibodies diluted in PB 0.1M for 1h at room temperature, then rinsed in PB 0.1M 3 times for 5 minutes each, and finally incubated 7 minutes with DAPI (1:5000 in PB 0.1 M) for nuclear staining. After washing, sections were mounted on glass slides and coverslipped under Mowiol medium.

#### Digital image acquisition

Images were acquired with a Zeiss Axio Imager Z2 microscope (Zeiss, Germany). Alexa 488 was imaged using a 495 nm beam splitter with an excitation wavelength set at 450/490 nm and an emission wavelength set at 500/550 nm. Alexa 568 was imaged using a 570 nm beam splitter with an excitation wavelength set at 538/562 nm and an emission wavelength set at 570/640 nm. Alexa 647 was imaged by using a 660 nm beam splitter with an excitation wavelength set at 625/655 nm and an emission wavelength set at 665/715 nm. Nuclear staining (Hoechst) was imaged by using a 395 nm beam splitter with an excitation wavelength set at 335/383 nm and an emission wavelength set at 420/470 nm. Images were acquired with a Plan-Apochromat 20X objective (numerical aperture NA = 0.80; thread type M27). Z-stack images were taken every 1 μm, for a total of 20 μm per sections. All the images are presented as maximal intensity projections (MIP) of three-dimensional volumes along the optical axis. Images to be used for figures were adjusted for brightness and contrast using Photoshop (Adobe Systems, San Jose, CA).

#### Cell counting

The quantification of nNOS neurons co-expressing p-nNOS was undertaken by counting the number of single-labeled nNOS-positive neurons and double-labeled nNOS/p-nNOS in the OV, represented respectively by plate 50 of the Allen Mouse Brain Atlas. Cell counts were carried out unilaterally, represented as percentage of nNOS neurons co-expressing p-nNOS, and the values from each mouse were averaged to determine mean counts and SEM for each age group.

### LH ELISA

Blood sampling to measure the circulating LH concentration was collected from the tale for pulsatility test. Diestrus female mice were gently restrained, and tail-tip blood sampling was taken at 10 min intervals. 4 μL of whole blood were collected into Eppendorf tubes pre-loaded with 46 μL of 0.01M PBS-0.05% Tween, thoroughly mixed, and placed on ice until storage at −80 °C. LH concentration was measured using a highly sensitive Enzyme-linked Immunosorbent Assay (ELISA) recently described in Kreisman et al., 2022. Briefly, we used a 96-well high-affinity binding microplate (Corning) coated with 50 μL of primary capture antibody (bovine LHβ subunit, 518B7; L. Sibley; University of California, UC Davis) diluted at 1µg/ml in 0.01M PBS. The mouse LH-RP was provided by Dr. Albert F. Parlow (National Hormone and Pituitary Program, Torrance, California, USA) and used to generate a standard curve with a twofold serial dilution. The detection primary antibody (Mouse Monoclonal LH antibody cat: Medix, 5303 SPRN SPRN-5) was biotinylated using the EZ-Link NHS-PEG4 Biotinylation Kit (Thermo Scientific, catalog No. PI21455) and used at a concentration of 1µg/ml, diluted in in 0.01M PBS-0.05%Tween, 5 % kim milk powder. For the final step, the poly-HRP Streptavidin (Thermo Fisher, Cat# N200) was diluted at 1:8000 in 0.01M PBS-0.05% Tween, then revealed with 100 μL of 1-Step Ultra TMB-Elisa Substrate Solution (ThermoFisher Scientific, cat. #34028). The colorimetric reaction was stopped with stop solution of 50 μL of 3M HCl. Optical density (OD) was read at 490 nm, the 650 for background subtraction.

### ELECTROPHYSIOLOGY

#### Brain slice preparation

Electrophysiological recordings were performed on living brain slices from 8 to 12-week-old mice. Mice were put under isoflurane anesthesia and killed by decapitation. The brain was dissected and rapidly placed in ice-cold aCSF containing: 120 mM NaCl, 3.2 mM KCl, 1 mM NaH_2_PO_4_, 26 mM NaHCO_3_, 1 mM MgCl_2_, 2 mM CaCl_2_, 10 mM glucose (300 mOsm, pH 7.4) and bubbled with 95% O_2_ to 5% CO_2_. 200 µm coronal slices containing the preoptic area were cut using a VT1200 vibratome (Leica). Slices were incubated at 34°C in oxygenated aCSF for a recovery period of 1 hour, and then placed at room temperature until patch-clamp recording.

#### Patch-clamp recording

Individual brain slices were placed in a submerged recording chamber (Warner Instruments) and continuously perfused at a rate of 2-3mL/min with oxygenated aCSF maintained at 32.8°C by a heater controller (TC-344C-Warner Instrument). GnRH neurons were visualized under x10 and x40 magnification using an upright fluorescence microscope with infrared differential interference contrast (DM-LFSA, Leica) and an ORCA-Frash4.0 digital CMOS camera (Hamamatsu). Recording pipettes were pulled from borosilicate glass capillaries (1.5 mm outer diameter, 1.12 mm inner diameter; World Precision Instruments) using a P1000 Flaming Brown puller (Sutter Instruments) and have a resistance of 7 MΩ when filled with an internal solution containing 140 mM K-gluconate, 10 mM KCl, 1 mM EGTA, 2 mM Na_2_-ATP and 10 mM HEPES (290 mOsm, pH 7.3 with KOH). Whole-cell patch-clamp recordings were performed in current-clamp mode using a Multiclamp700B Amplifier, digitized with the Digidata 1322A interface and acquired with pClamp 10.2 software (Molecular Devices). All drugs tester (Kisspeptin 10 nM, L-NAME 1 mM, DEA/NO 100 µM) were applied to the perfusion system after stable baseline recording. Recordings were analyzed using Clampfit 10.2 pClamp software (Molecular Devices). For each recording, the membrane potential and mean firing rate were determined before and during the bath application of drugs. Neurons were considered to have responded if there was a > 20% change in firing rate during the drug application. Only cells which showed less than 20% change in access resistance throughout the recording were included in this study. Junction potential was corrected in the data analysis.

## ACKNOWLEDGMENTS

This work has been supported by the European Union Horizon 2020 research and innovation program No 847941 miniNO, the European Research Council (ERC) Synergy grant No 810331 WATCH, the DistAlz (ANR-11-LABEX-0009), European Genomic Institute for Diabetes (EGID, ANR-10-LABX-0046), the I-SITE ULNE (ANR-16-IDEX-0004), the Société Française d’étude de la Fertilité (SFEF) (MSc fellowship to J.D.) and the University of Lille (PhD fellowship to V.D.). The authors thank Meryem Tardivel and Antonino Bongiovanni (imaging core facility, BiCeL) and Julien Devassine (animal house) of the UMS2014-US41 for her expert technical assistance. Graphics were created with BioRender.com.

