## Supplementary material for "The preoptic Kisspeptin/nNOS/GnRH (KiNG) neuronal network regulates rhythmic LH release through a dual activation-inhibition mechanism": Supplemetary figures 1-5

### SUPPLEMENTARY FIGURES

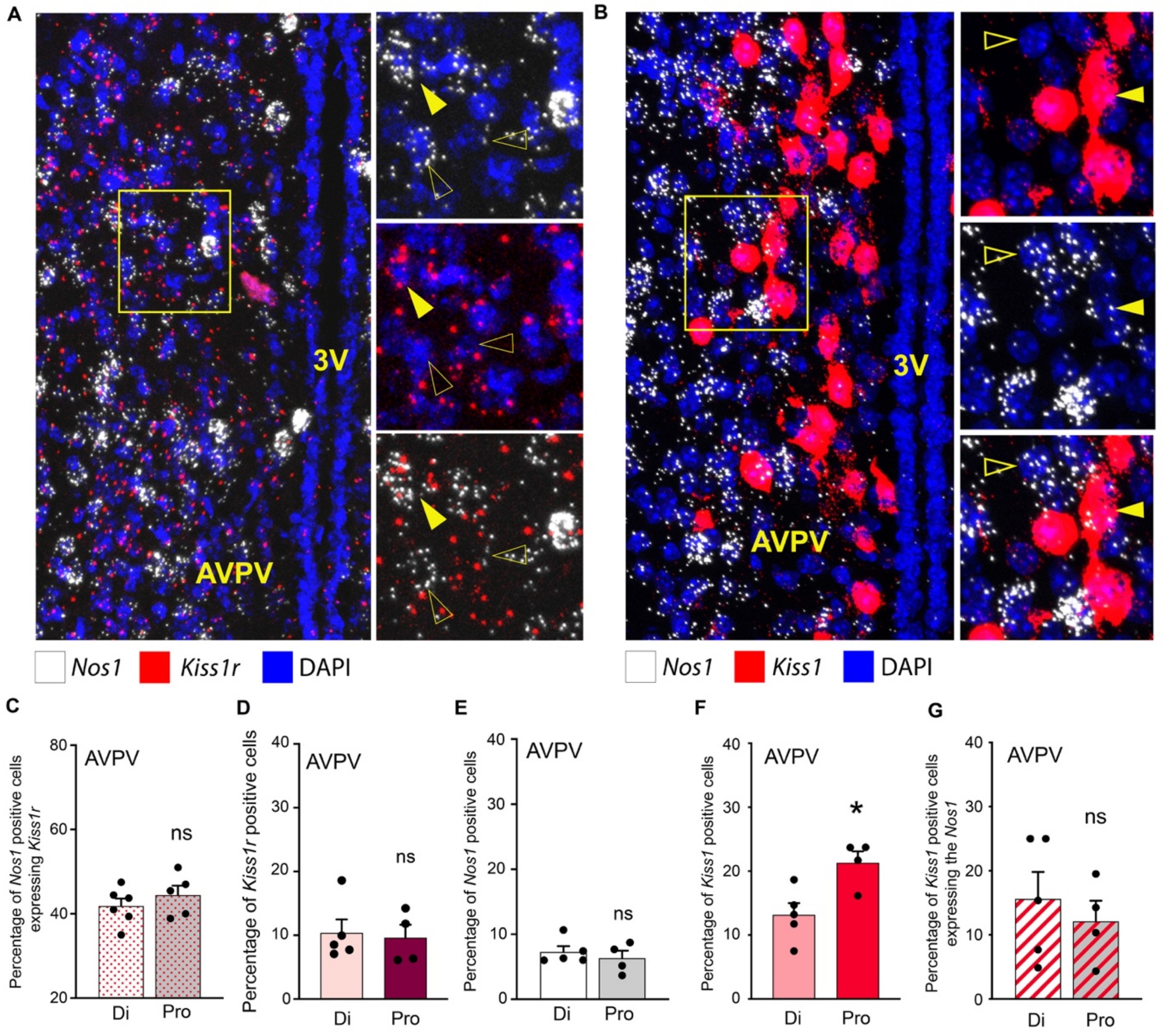


**Suppl. Figure 1. Kisspeptin-nNOS interaction dynamics in the AVPV of the cycling female mouse.** (A-B) Confocal florescent RNAscope revealing the anatomical interaction and differential expression of (A) *Nos1* (white), *Kiss1r* (red) or (B) *Nos1* (white) and *Kiss1* (red) mRNA in the female mouse preoptic area, in the organum vasculosum laminae terminalis (OV) and median preoptic nucleus (MePO). Magnifications on the right side of the illustrations indicate representative examples of *Nos1* mRNA positive cells co-expressing the *Kiss1r* or *Kiss1* mRNA (yellow arrowheads), as well as *Nos1*-negative cells (empty arrowheads). (C) Quantification of the percentage of cells positive for the *Nos1* mRNA also being positive for the *Kiss1r* mRNA in Diestrous and Proestrous in the region of the AVPV (Mann-Whitney t-test; Di , N=4 ; Pro N=5). (D-F) Quantification of the percentage of the cells (over total DAPI) positive for the (D) *Kiss1r*, (E) *Nos1* and (D) *Kiss1* mRNA in Diestrous and Proestrous in the region of the AVPV (Mann-Whitney; Di, N=5 ; Pro N=4, p=0.03). (G) Quantification of the percentage of cells positive for the *Kiss1* mRNA also being positive for the *Nos1* mRNA in Diestrous and Proestrous in the region of the AVPV (Mann-Whitney t-test; Di, N=5 ; Pro N=4). ns: non-significant, * P < 0.05. 3V; third ventricle; AVPV, anteroventral periventricular nucleus; Diestrous, Di; Proestous, Pro.

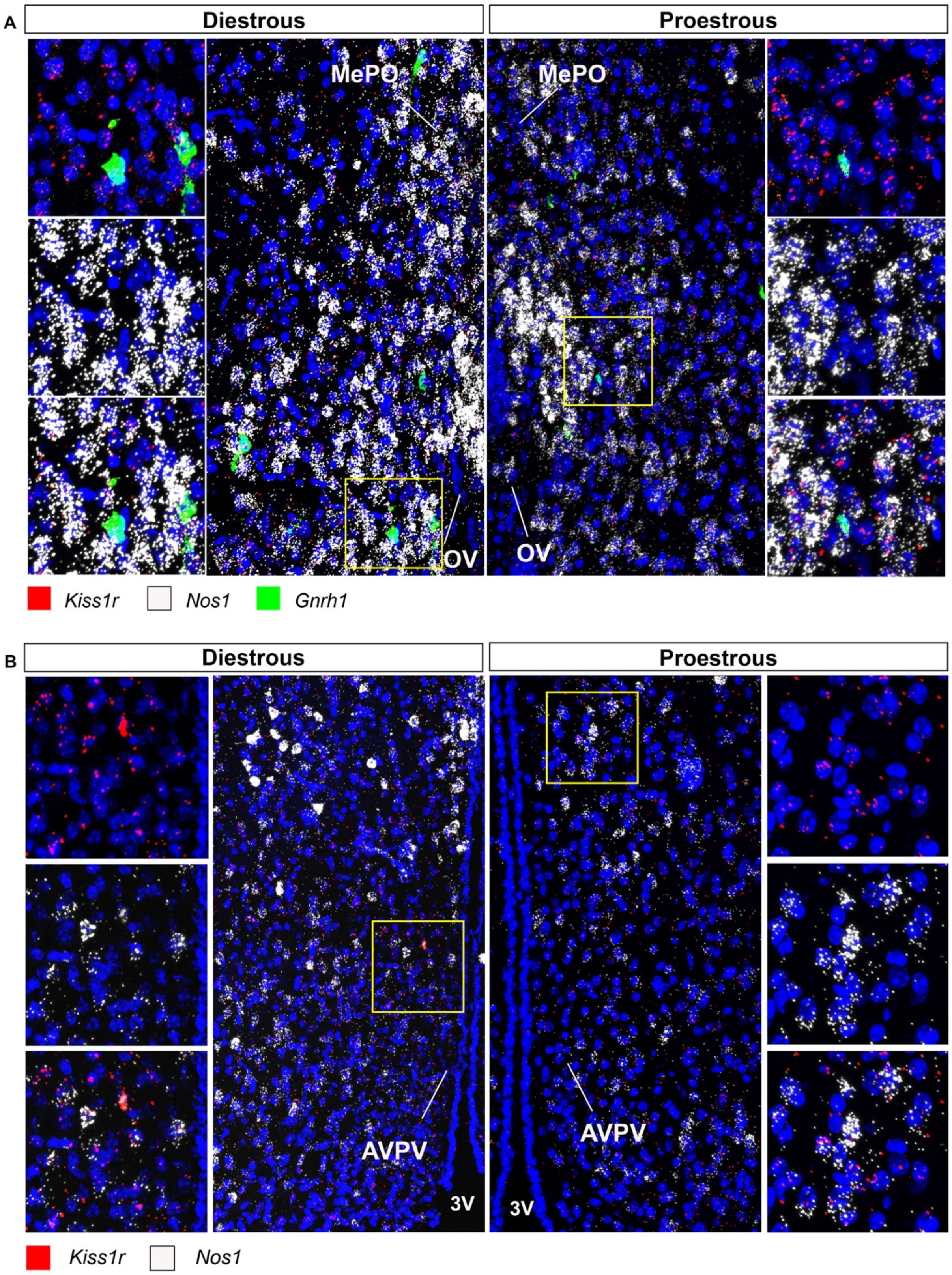


**Suppl. Figure 2. Kisspeptin receptor-nNOS interaction in the OV/MePO and AVPV of the cycling female mouse.**

(A) Confocal fluorescent RNAscope revealing the anatomical interaction and differential expression of *Nos1* (white), *Kiss1r* (red) and *Gnrh1* (green) mRNA in the female mouse preoptic area in the organum vasculosum laminae terminalis (OV) and median preoptic nucleus (MePO), during the stages of Diestrous (left) or Proestrous (right). Magnifications on the left and right side of the illustrations indicate representative examples of *Nos1* and *Gnrh1* mRNA positive cells co-expressing the *Kiss1r* mRNA. (B) Confocal fluorescent RNAscope revealing the anatomical interaction and differential expression of *Nos1* (white) and *Kiss1r* (red) mRNA in the female mouse anteroventral periventricular nucleus (AVPV) during the stages of Diestrous (left) or Proestrous (right). Magnifications on the left and right side of the illustrations indicate representative examples of *Nos1* mRNA positive cells co-expressing the *Kiss1r* mRNA. 3V, third ventricle; OV, organum vasculosum laminae terminalis; MePO, median preoptic nucleus; AVPV, anteroventral periventricular nucleus; Diestrous, Di; Proestous, Pro.

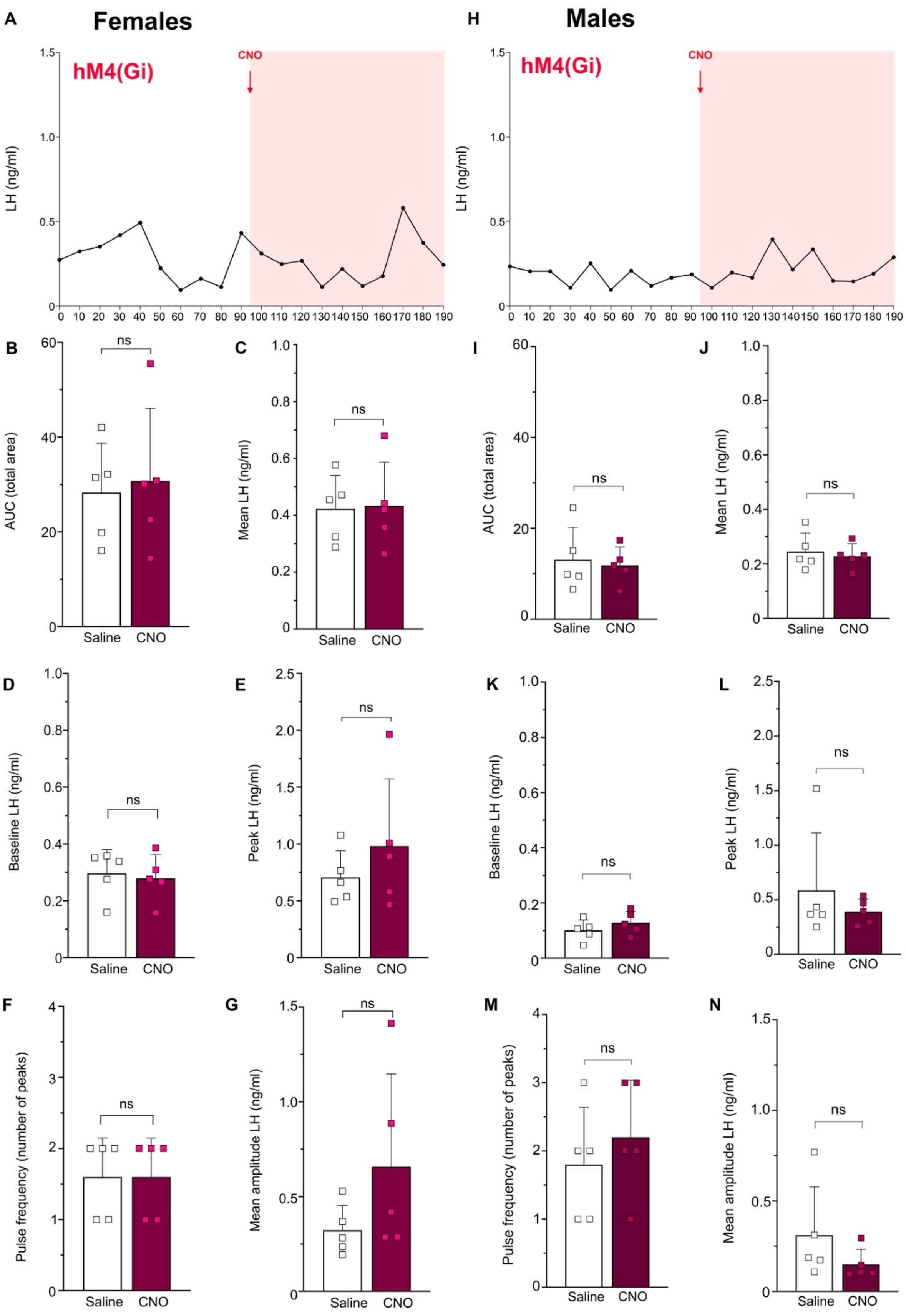


**Suppl. Figure 3. Effect of local inhibition of the nNOS $^{OV/MePO}$ neurons on the pulsatile LH release in female and male mice.** Representative LH pulse profiles and subsequent analysis of intact Nos1cre female (A-G) and male (H-N) mice injected into the MePO with the AAV-hSyn-DIO-hM4(Gi)-

mCherry. (B, I) Area Under the Curve (AUC) analysis, (C, J) Mean total LH, (D, K) Mean baseline LH levels, (E, L) Peak LH response, (F, M) Pulse frequency and (G, N) Mean amplitude of LH responses before (white) or after (purple) CNO administration (95$^{th}$ minute). N=5. ns: non-significant, $P > 0.05$.

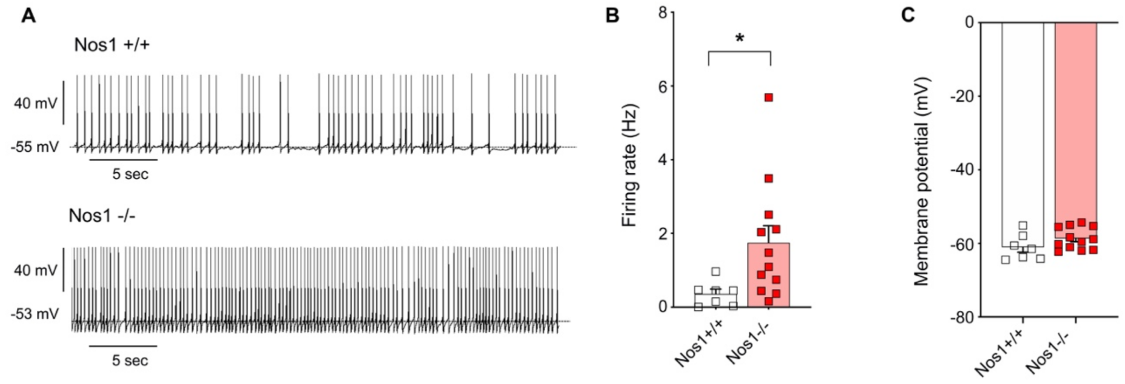


**Suppl. Figure 4. Effect of genetic ablation of nNOS activity on the GnRH neuronal activity in adult female mice in Diestrous.**

(A) Electrophysiological recordings of the spontaneous activity of preoptic area GnRH neurons in adult diestrous *Gnrh::Gfp; Nos1*$^{+/+}$ and *Gnrh::Gfp*;*Nos1*$^{-/-}$ bigenic mice. (B) *Gnrh::Gfp; Nos1*$^{+/+}$ firing rate and (C) membrane potential values are compared to those of *Gnrh::Gfp; Nos1*$^{-/-}$ mice (unpaired t-test; *Gnrh::Gfp; Nos1*$^{+/+}$ , n=7 cells, N=6 mice ; *Gnrh::Gfp*;*Nos1*$^{-/-}$, n=12 cells, N=6 mice). * $P < 0.05$. Values indicate means ± SEM.

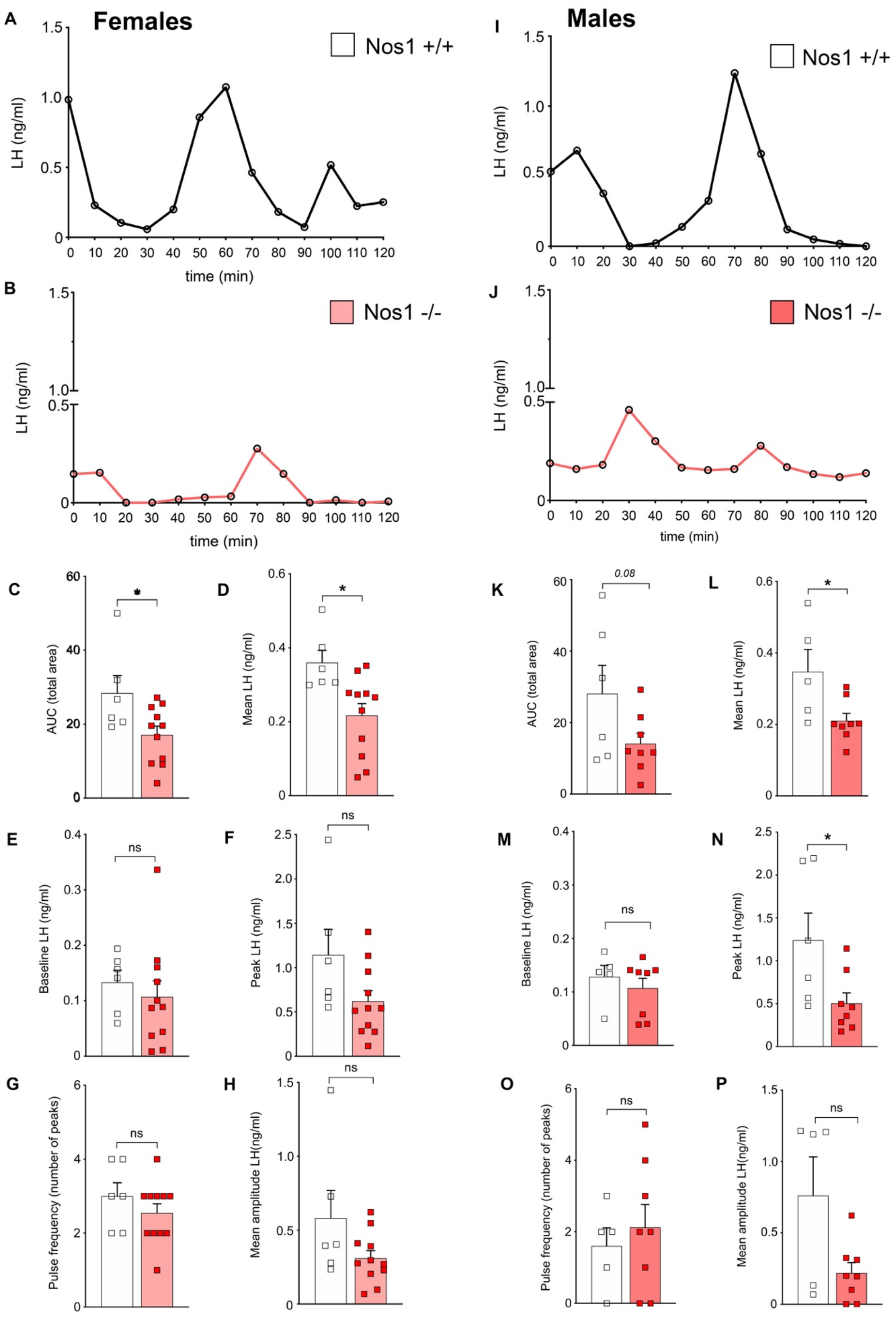


**Suppl. Figure 5. Effect of nNOS genetic deficiency on the LH pulsatile release in female and male mice.** (A-H) Representative LH pulse profiles of an intact Nos1 +/+ (A) and Nos1 -/- (B) female mouse during diestrous. (B) Area Under the Curve (AUC) analysis (unpaired t-test, p= 0.02, n=6, 11), (C) Mean total LH (unpaired t-test, p= 0.01, n=6, 11), (D) Mean baseline LH levels, (E) Peak LH response, (F) Pulse frequency and (G) Mean amplitude of LH responses in intact Nos1 +/+ (white) and Nos1 -/- (red) females at diestrous. N=. (I-P) Representative LH pulse profiles of an intact Nos1 +/+ (A) and Nos1 -/- (B) male mouse (B) Area Under the Curve (AUC) analysis (unpaired t-test, p= 0.08, n=6, 8), (C) Mean total LH (unpaired t-test, p= 0.03, n=5, 8), (D) Mean baseline LH levels, (E) Peak LH response (unpaired t-test, p= 0.03, n=6, 8), (F) Pulse frequency and (G) Mean amplitude of LH responses in intact Nos1 +/+ (white) and Nos1 -/- (red) males. ns: non-significant, * $P < 0.05$. Values indicate means ± SEM.
